# TBK1 interacts with tau and enhances neurodegeneration in tauopathy

**DOI:** 10.1101/2020.06.17.157552

**Authors:** Measho H. Abreha, Shamsideen Ojelade, Eric B. Dammer, Zachary T. McEachin, Duc M. Duong, Marla Gearing, Gary J. Bassell, James J. Lah, Allan I. Levey, Joshua M. Shulman, Nicholas T. Seyfried

**Affiliations:** Department of Biochemistry, Emory University School of Medicine, Atlanta, GA, 30322, USA; Department of Neurology, Emory University School of Medicine, Atlanta, GA, 30322, USA; Department of Cell Biology, Emory University School of Medicine, Atlanta, GA, 30322, USA; Department of Pathology and Laboratory Medicine, Emory University School of Medicine, Atlanta, GA, 30322, USA; Center for Neurodegenerative Diseases, Emory University School of Medicine, Atlanta, GA, 30322, USA; Department of Molecular and Human Genetics, Baylor College of Medicine, Houston, TX 77030, USA; Department of Neuroscience, Baylor College of Medicine, Houston, TX 77030, USA; Department of Neurology, Baylor College of Medicine, Houston, TX 77030, USA; Jan and Dan Duncan Neurological Research Institute, Texas Children’s Hospital, Houston, TX 77030, USA

**Keywords:** TANK-binding kinase 1 (TBK1), microtubule associated protein tau (MAPT), neurofibrillary tangles (NFTs), Alzheimer’s disease (AD), familial frontotemporal dementia and parkinsonism linked to chromosome 17 (FTDP-17), co-immunoprecipitation (co-IP), mass spectrometry (MS), post-translational modification (PTM), *Drosophila*, IκB kinase (IKK)

## Abstract

One of the defining pathological features of Alzheimer’s Disease (AD) is the deposition of neurofibrillary tangles (NFTs) composed of hyperphosphorylated tau in the brain. Aberrant activation of kinases in AD has been suggested to enhance phosphorylation and toxicity of tau, making the responsible tau-directed kinases attractive therapeutic targets. The full complement of tau interacting kinases in AD brain and their activity in disease remains incompletely defined. Here, immunoaffinity enrichment coupled with mass spectrometry (MS) identified TANK-binding kinase 1 (TBK1) as a tau-interacting partner in human AD cortical brain tissues. We validated this interaction in both human AD and familial frontotemporal dementia and parkinsonism linked to chromosome 17 (FTDP-17) caused by mutations in *MAPT* (R406W) postmortem brain tissues as well as human cell lines. Further, we document increased TBK1 activity in both AD and FTDP-17 and map the predominant TBK1 phosphorylation sites on tau based on *in vitro* kinase assays coupled to MS. Lastly, in a *Drosophila* tauopathy model, activating expression of a conserved TBK1 ortholog triggers tau hyperphosphorylation and enhanced neurodegeneration, whereas knockdown had the reciprocal effect, suppressing tau toxicity. Collectively, our findings suggest that increased TBK1 activity may promote tau hyperphosphorylation and neuronal loss in AD and related tauopathies.

## Introduction

Deposition of extracellular amyloid-β (Aβ) plaque and accumulation of intracellular neurofibrillary tangles (NFTs) composed of the microtubule binding protein tau in the brain are the major hallmarks of Alzheimer’s Disease (AD) (1-3). The degree of NFT burden in the brain is closely associated with synaptic loss and cognitive decline, therefore, making tau a major therapeutic target (4,5). Tau is a phosphoprotein that is required for binding and stabilization of microtubules under physiological conditions (6). In AD, tau undergoes hyperphosphorylation among other post-translational modifications (PTMs), leading to accumulation of NFTs (7-9). Tau phosphorylation is regulated by tau-directed kinases and phosphatases the aberrant levels and activity of which have been suggested to impact tau hyperphosphorylation and aggregation (10,11). Towards this end, the identification of new tau-directed kinases could reveal potential targets and contribute to further development of therapeutic drugs in the treatment of AD.

Mass spectrometry (MS) based proteomics has emerged as a powerful approach to define global changes in protein abundance and PTMs linked to disease mechanisms (12). MS-based proteomics can also be combined with affinity enrichment strategies, such as immunoprecipitation (IP), to identify interacting partners of key proteins linked to disease pathogenesis (13,14). We recently coupled co-immunoprecipitation (co-IP) with MS to identify and quantify tau interacting partners (i.e., tau interactome) from control and AD brain tissues, which led to the identification of over 500 proteins that are enriched specifically with tau in AD (15). How these interacting proteins relate to pathologic tau phosphorylation remains to be determined.

From these tau co-IP datasets in AD brain (15), we mapped tau phosphorylation sites and 21 total kinases, of which 18 have been previously described to interact or directly phosphorylate tau (16,17). Of the three novel tau interacting kinases identified, we further validated TBK1, a pleiotropic serine threonine kinase belonging to the non-canonical IKK family of kinases (29,30), best characterized for its role in innate immunity signaling (31-35), selective autophagy pathways (36-38), energy metabolism (39,40), tumorigenesis (41), and microtubule dynamics (42). Notably, genetic studies have also identified mutations in TBK1 gene as causal for neurodegenerative diseases including frontotemporal dementia (FTD) and amyotrophic lateral sclerosis (ALS) (43-47). However, a role for TBK1 in modifying tau phosphorylation and toxicity in AD and other tauopathies is not known.

Here we show increased TBK1 activity in both AD and frontotemporal dementia and parkinsonism linked to chromosome 17 (FTDP-17) and map the predominant TBK1 phosphorylation sites on tau based on *in vitro* kinase assays coupled to MS. TBK1 over-expression studies in human cells and *Drosophila* neurons further confirmed the role of TBK1 activation in tau hyperphosphorylation in model systems. Finally, TBK1 overexpression and knockdown in *Drosophila* can reciprocally increase and decrease tau-induced neurodegeneration. Together these data support a hypothesis that TBK1 activity may enhance tau phosphorylation and neuronal loss in AD and related tauopathies.

## Results

### Identification of tau phosphorylation sites and interacting kinases in AD brain

Tau hyperphosphorylation highly correlates with degree of synaptic loss and cognitive decline (4,5). In our recently published MS dataset based on immunoaffinity enriched tau from AD (n=4) and control (n=4) postmortem tissues (15) (prefrontal cortex), we identified increased intensities of tau phosphopeptides, consistent with the expected tau hyperphosphorylation in AD (**Figure 1A and Table S1**). Tau directed kinases tightly regulate tau phosphorylation and their aberrant activity has been previously reported in AD (11,48). Moreover, Gene Ontology (GO) analysis (15) revealed significant enrichment of terms related to kinase activity (e.g. “GTPase activity”, “kinase signaling”, and “kinase binding”) among proteins showing increased association with tau in AD (**Figure 1B**). Further analysis of tau interacting partners enriched in AD identified 21 kinases including several well-established tau-directed kinases such as glycogen synthase kinase 3β (GSK3β) (11), cyclin dependent kinase 5 (CDK5) (49), mitogen-activated protein kinases (MAPK1 and MAPK15) (17), calcium/calmodulin dependent protein kinases (CAMK2A, CAMK2B, CAMK2D and CAMK2G) (50), and TAOK1 (51). Three previously unreported putative novel tau-directed kinases (TNIK, DAPK3 and TBK1) were also identified (**Figure 1C and Figure S1**). Notably, TBK1 is a serine threonine kinase belonging to the non-canonical IKK family of kinases with roles in the innate immunity signaling (29,31) and autophagy pathway (34,52). We focused our attention on TBK1, as recent studies have identified mutations in TBK1 gene as causal for neurodegenerative diseases including FTD and ALS (44,46). However, a role for TBK1 in modifying tau phosphorylation and toxicity in AD and related tauopathies has not been established.

**Figure 1.**
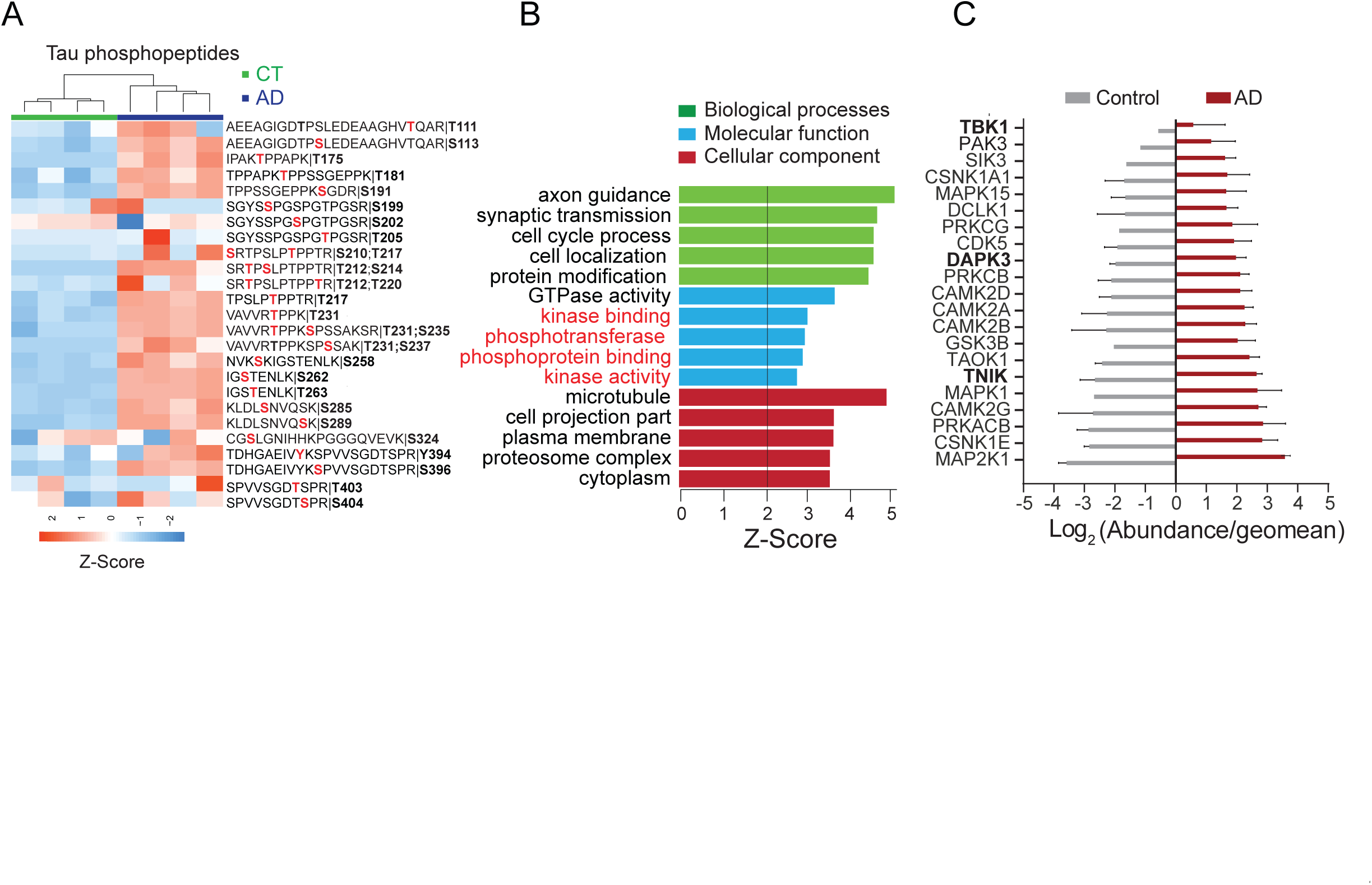
Identification of tau phosphorylation and interacting kinases by co-IP and MS analysis of human postmortem brains. A) Hierarchical clustering of 26 tau phosphopeptide intensities identified from tau co-IP analysis using control and AD postmortem brain tissues (*n*=4, each) shows increased tau phosphorylation on 20 sites in AD. B) Gene ontology (GO) analysis of AD tau interacting proteins (n=513) identified over representation of proteins associated with kinase activity. C) Histogram displaying log_2_ (abundance/geomean) reveals 21 Tau interacting kinases. Novel Tau interacting kinases (TBK1, DAPK3 and TNIK) are highlighted in bold.

### Increased tau interaction and activity of TBK1 in AD and FTDP-17 brain

To validate our proteomics findings, we performed co-IP of tau from AD and control (n=2, each) postmortem cortical brain lysates using a tau monoclonal antibody followed by western blot analysis for TBK1. Analysis of the brain samples by western blot showed a high molecular weight (HMW) tau species in AD samples reflective of oligomeric and modified tau isoforms in disease (53,54) (**Figure 2A**). Assessment of TBK1 in the brain samples (1x lysates) revealed similar levels of TBK1 in control and AD samples indicating that steady-state levels of the kinase are unchanged in disease (**Figure 2A, lanes 1 & 2, versus lanes 5 & 6**). In contrast, western blot analysis of tau immunoprecipitates showed strong bands for TBK1 (∼100kDa) in AD compared to controls, indicating a preferential interaction between TBK1 and AD tau (**Figure 2B**). To determine whether the increased TBK1 co-IP in AD is simply due to increased tau abundance in AD, we doubled the total protein amount in controls (2x lysate), which, as expected, revealed increased levels (∼2-fold) of tau and TBK1 in the inputs (**Figure 2A, lanes 3 & 4**). However, despite doubling the tau and TBK1 levels we did not observe TBK1 interactions with tau immunoprecipitates in control brain (**Figure 2B, lanes 3 & 4**). This suggests that TBK1 has increased affinity for AD tau due to potential phosphorylation and/or conformational changes.

**Figure 2.**
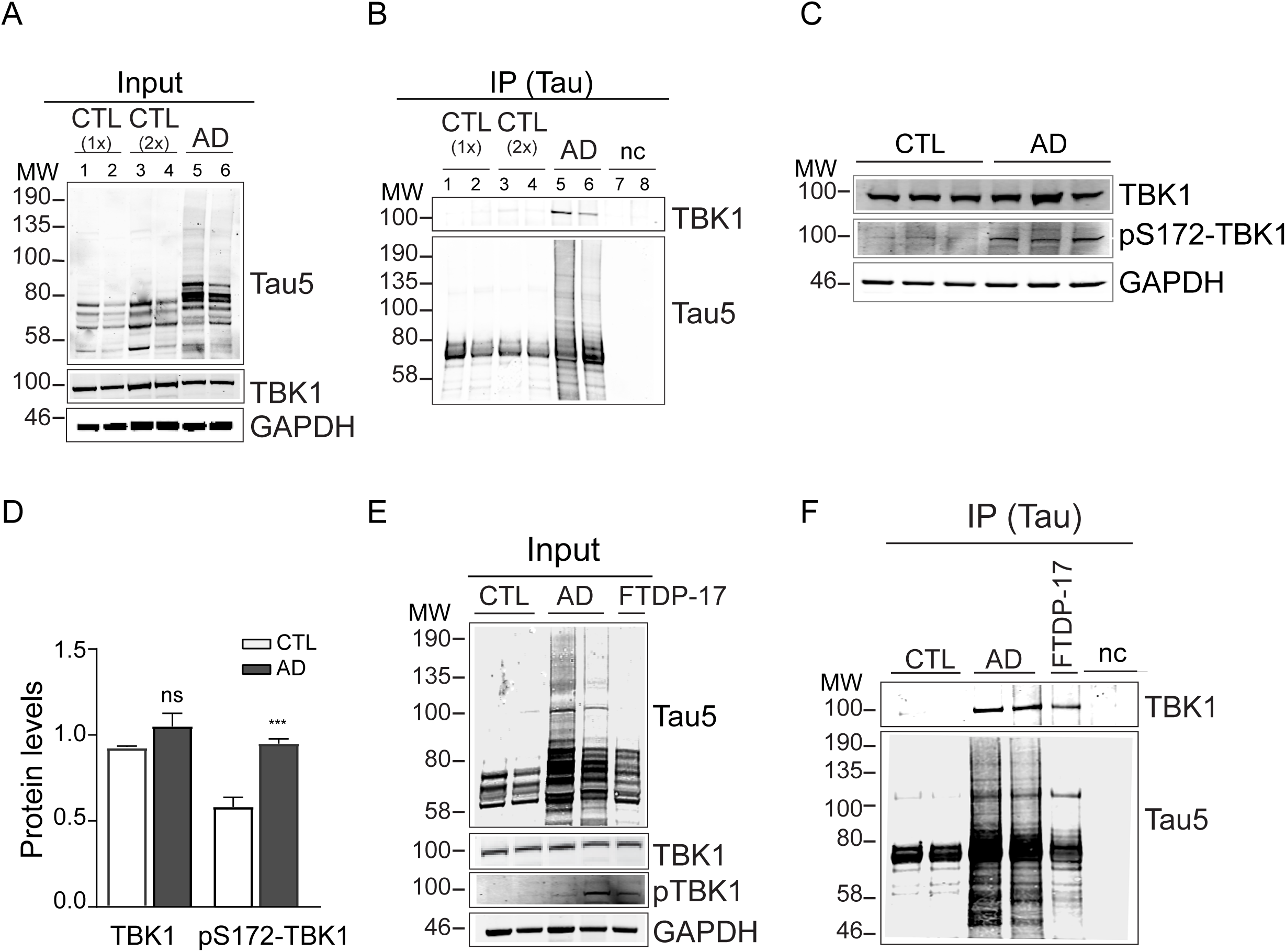
Increased tau interaction and activity of TBK1 in AD and FTDP-17 (R406W) brains. A) Western blot analysis of two AD and control postmortem brains (input) using Tau5 and TBK1 antibodies showed high molecular weight (HMW) tau species in AD and equal levels of TBK1 across AD and control (CTL 1x) cases. Doubling the input (2x CTL) shows increased levels of both tau and TBK1. B) IP of tau followed by western blot for TBK1 shows a band in AD samples compared to CTL 1x and 2x samples. Lysates incubated with beads alone were used as a negative control (nc) for both AD and control lysates. C) Western blot analysis of postmortem brain tissues from control and AD cases (n=3, each) for total TBK1 and active TBK1 (pS172-TBK1) showed increased pS172-TBK1 levels in AD compared to controls. D) Quantification of the pS172-TBK1 levels in AD and control samples shows a significant increase in AD (T-test, *** = *P*<0.001). E) Western blot analysis of control (*n*=2), AD (*n*=2) and a FTDP-17 (R406W) postmortem brain samples (input) using total TBK1 and active TBK1 (pS172-TBK1) showed increased TBK1 activation in AD and FTDP-17 (R406W) while TBK1 remains the same. E) Co-IP for tau followed by western blot for TBK1 shows TBK1 interaction with AD and FTDP-17 tau. GAPDH was used a loading control. Lysates incubated with beads alone were used as negative controls (nc).

Given that TBK1 interacts with AD tau, yet steady state TBK1 levels remain unchanged, we hypothesized that TBK1 activation is elevated in AD brain. Notably, TBK1 activation is induced by trans-autophosphorylation of serine 172 (S172) in the activation loop within the kinase domain (55,56). Therefore, to measure activated TBK1 levels, we performed western blot analysis of AD and control postmortem brain lysates with a pS172-TBK1 antibody. We observed an increase in pS172-TBK1 levels in AD brain samples (**Figure 2C**) and quantification of both total TBK1 and active TBK1 levels shows a statistically significant (p = 0.0056) increase in active TBK1 in AD (**Figure 2D**). Thus, although TBK1 steady state levels remain unchanged in AD, our data indicate that TBK1 activation and interaction with AD tau is increased.

To assess whether the increased AD tau and TBK1 interaction was unique to AD or shared with other tauopathies, we performed co-IP for tau in brain cortical tissues from a FTDP-17 case caused by a missense mutation (R406W) within the C-terminus of MAPT. Notably, FTDP-17 cases with this particular mutation exhibit an AD-like clinical phenotype including relatively late onset amnestic dementia and longer disease duration as well as accumulation of hyperphosphorylated tau in NFTs (57). Like AD, the FTDP-17 (R406W) case showed increased HMW tau species and TBK1 activation (**Figure 2E**). Furthermore, tau showed increased interaction with TBK1 in FTDP-17, but not control cases (**Figure 2F**). Overall, our data suggest that increased activation and interaction of TBK1 with tau may be common across these tauopathies.

### TBK1 directly phosphorylates tau

We next sought to determine whether TBK1 directly phosphorylates tau. To identify putative sites of TBK1 directed phosphorylation of tau, we performed *in vitro* kinase assays, whereby recombinant purified tau protein was incubated in kinase buffer with ATP in the presence or absence of recombinant active TBK1. Following tryptic digestion and MS analysis of the *in vitro* reactions, we identified phosphopeptides corresponding to 9 phosphorylation sites on tau (T30, S191, S198, S214, S285, S289, S305, S324, and S356) (**Table S2**). These sites are distributed across the N-terminal acidic domain (T30), proline-rich mid-domain (S191, S198, S214), and microtubule binding repeat domains (S285, S289, S305, S324, S356) of tau (**Figure 3A and B**). Tau S214 phosphorylation is indicated to enhance microtubule dissociation and correlates with NFTs formation in AD (58). Of note, S214 phosphorylation is part of the AD associated AT100 tau epitope and is described to be a priming/pre-phosphorylation signal for subsequent tau phosphorylation by GSK3β (18). Thus, given the biological importance and strong increase in disease (described below) we chose S214 tau phosphorylation as a surrogate for TBK1-directed tau phosphorylation. Western blot analysis using a pS214-tau antibody showed loss of pS214-tau signal in the absence of ATP or the presence of 100 μM or 200 μM of a selective TBK1 inhibitor, BX795 (59,60), whereas active TBK1 (pS172-TBK1) and total tau levels remained unaltered in all conditions (**Figure 3C**). Thus, these *in vitro* analyses confirm that tau is a direct substrate of TBK1, and that it can be blocked by a small molecule TBK1 inhibitor.

**Figure 3.**
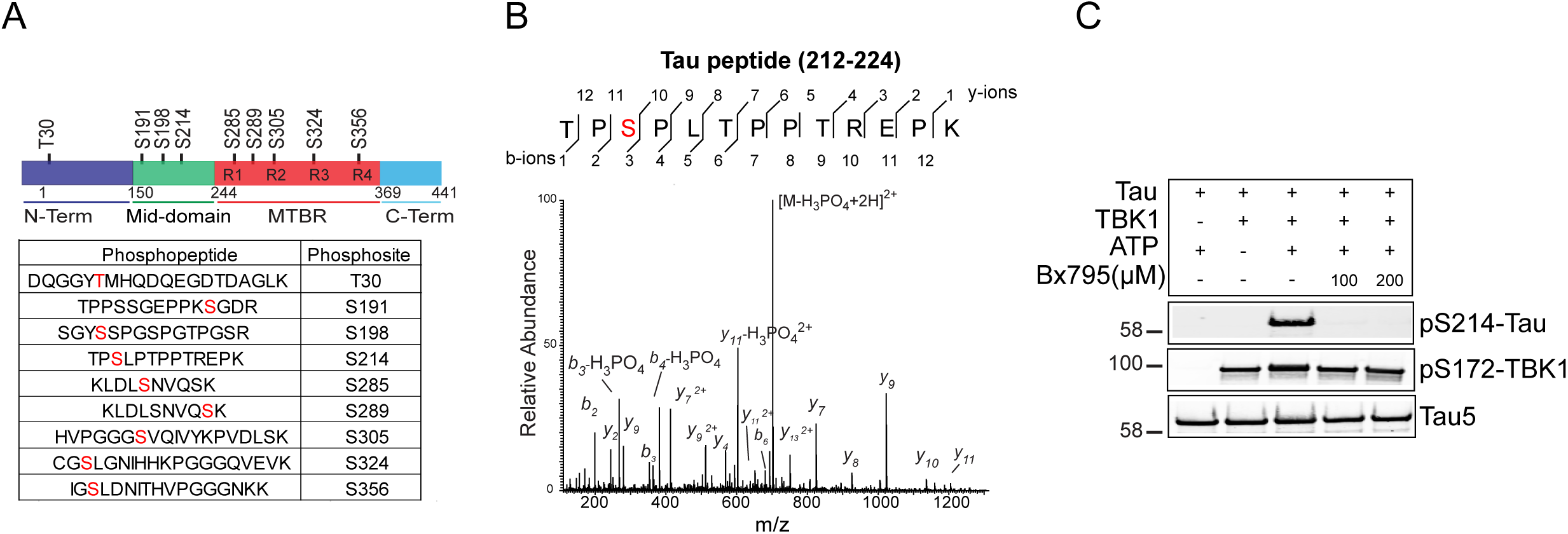
TBK1 directly phosphorylates tau. A) MS analysis of *in vitro* TBK1 kinase activity assay using recombinant tau and active TBK1 in the presence of ATP identifies TBK1 phosphorylation sites on tau. A schematic depicting TBK1-mediated phosphorylation sites across tau domains and a list of tau phosphopetides identified in the *in vitro* kinase assay. TBK1 targeted S/T residues are indicated in red. B) MS/MS spectrum of a tau phosphopeptide with S214 phosphorylation site (residues 212-224) shows b- and y-ions and the diagnostic neutral loss of phosphoric acid (−98 Da) on the precursor peptide. C) Western blot analysis of the TBK1 *in vitro* kinase activity assays using pS214-Tau antibody shows loss of S214 phosphorylation in the presence of a TBK1 inhibitor (BX795) or absence of ATP.

### Increased phosphorylation of TBK1-mediated tau phosphosites in AD and FTDP-17 brain

To assess the pathological significance of the *in vitro* TBK1-directed phosphosites mapped on tau, we analyzed tau phosphopeptides identified and quantified from tau IPs across control and AD cases (*n*=4, each) (**Figure 1A**). Comparison of brain tau phosphosites with the *in vitro* TBK1 sites showed a high degree of overlap in tau phosphosites (S191, S214, S285, S289, and S324), which were all elevated in AD brain (**Figure 4A and Figure 1A**). Among the tau phosphosites, we validated S214 phosphorylation, which showed robust signal in both AD and FTDP-17 cases by western blot (**Figure 4B and C**). Similarly, immunohistochemical staining of AD and FTDP-17 cases showed increased tau S214 phosphorylation and accumulation of S214 phosphorylated tau in NFTs (**Figure 4D**), further highlighting similarities between the tauopathies. Collectively our findings link TBK1 activation in AD and FTDP-17 brain to increased phosphorylation of tau on five sites, including S214.

**Figure 4.**
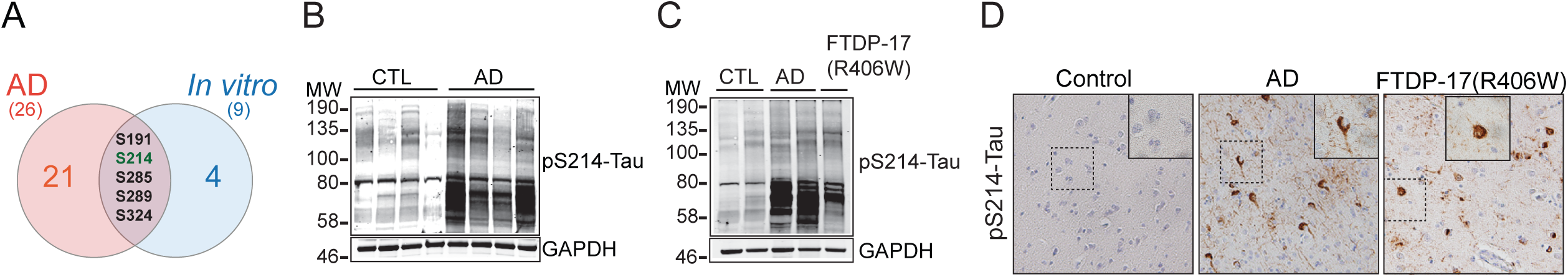
TBK1-directed tau phosphosites are increased in AD and FTDP-17 (R406W) brain tissue. A) A Venn diagram shows an overlap in tau phosphosites from the TBK1 *in vitro* kinase assay and AD brain tau IP. B) Increased tau pS214 phosphorylation in AD is confirmed using western blot analysis of AD and control postmortem brain samples (*n*=4, each). C) Western blot analysis using pS214-Tau antibody shows increased S214 phosphorylation in AD and FTDP-17 (R406W) tau compared to controls. GAPDH was used as a loading control D) Immunohistochemistry staining using pS214-Tau antibody shows increased tau S214 phosphorylation in AD and FTDP-17 (R406W) brain sections compared to controls.

### TBK1 alters tau phosphorylation in HEK-293 cell lines

To extend our *in vitro* findings, we sought to assess if TBK1 can interact with and phosphorylate tau in human cell lines. HEK-293 cell lines over-expressing either recombinant GFP-tau alone or both GFP-tau and recombinant Myc-TBK1 were used to assess TBK1-tau interactions (**Figure S2A**). IP of tau using a GFP antibody followed by western blotting showed strong association of tau with both total TBK1 (Myc-TBK1) and activated TBK1 (pS172-TBK1) (**Figure S2B**). Furthermore, reciprocal IP using a Myc antibody to enrich recombinant TBK1 also showed a strong association of TBK1 with tau (**Figure S2C**), which is consistent with the interaction observed in human AD and FTDP-17 brain (**Figure 2B and F**). To assess if TBK1 activation enhances tau phosphorylation in HEK-293 cells, we over-expressed GFP-tau and Myc-TBK1. Western blot of cell lysates using pS214-tau antibody showed a significant increase in tau phosphorylation at S214 (**Figure 5A and B**), consistent with in vitro kinase assays (**Figure 3C**). In contrast, over-expression of a specific phospho-null tau mutant (tau-null), in which the nine serine and threonine residues that can be phosphorylated by TBK1 were substituted by alanine (**Figure 3A**), resulted in a significant decrease in overall tau phosphorylation. supporting that TBK1-directed tau phosphorylation in cells is primarily directed at these nine residues (**Figure 5A and B**). To determine the effect of TBK1 inhibition on tau phosphorylation, HEK-293 cells were treated with a TBK1 inhibitor, BX795, followed by western blot analysis using GFP and pS214-tau antibodies to detect total and phosphorylated tau, respectively (**Figure 5C**). Following BX795 addition, the levels of activated TBK1 (pS172) and S214 tau phosphorylation significantly decreased (p<0.001), whereas the steady-state TBK1 levels remained the same, further supporting TBK1-directed tau phosphorylation in cells (**Figure 5D**). Overall, our data demonstrate that TBK1 can mediate tau phosphorylation in cells that are linked to pathologic sites in AD and related tauopathies.

**Figure 5.**
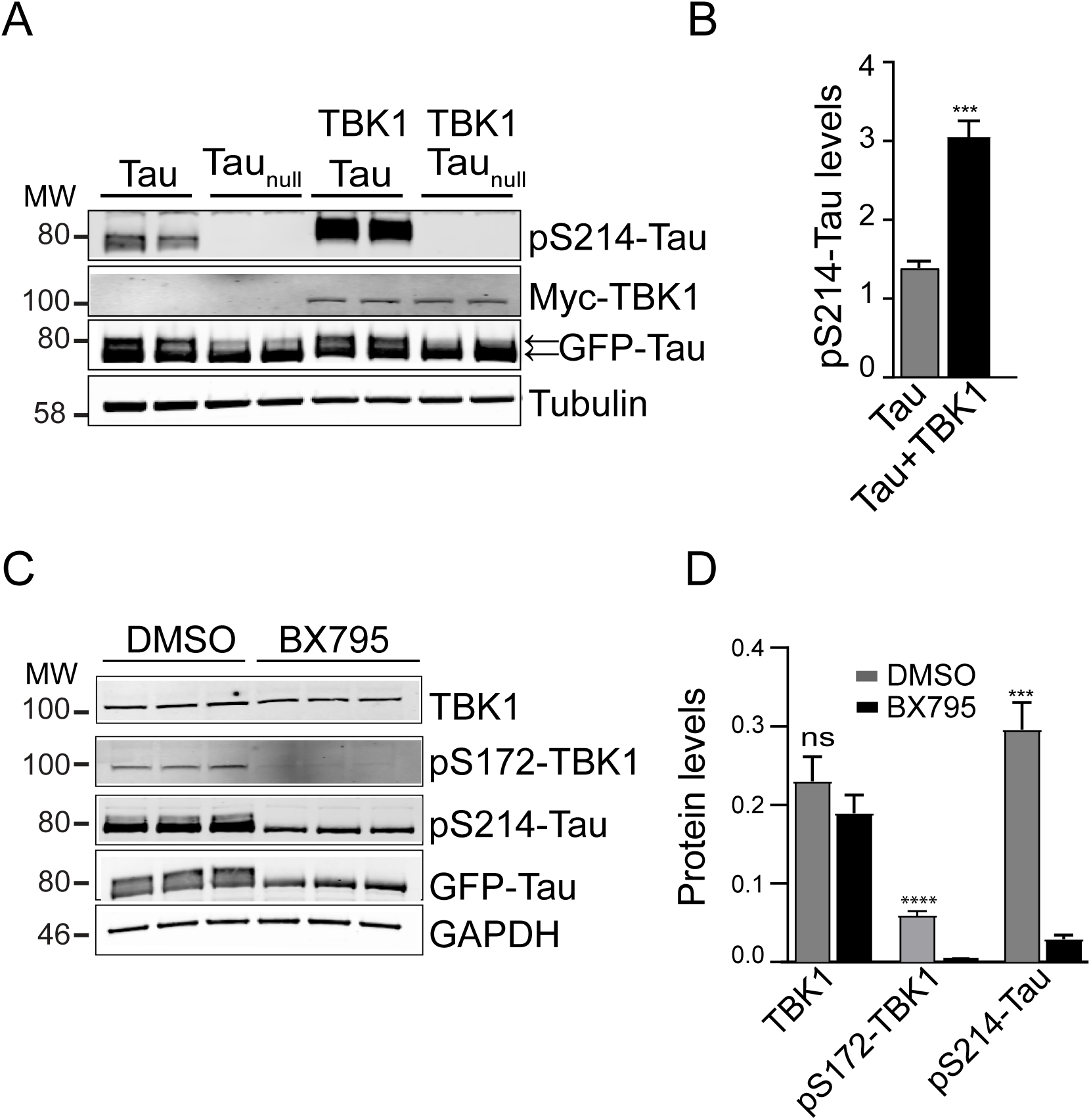
Over-expression or inhibition of TBK1 alters tau phosphorylation in HEK-293 cells. A) HEK-293 cells were transfected with recombinant myc-tagged TBK1 (Myc-TBK1) and either recombinant GFP-tagged tau (GFP-Tau) or a mutant tau (GFP-Tau_null_) with all nine TBK1 sites substituted with alanine. Cells were harvested 48 hrs. post-transfection and subjected to western blot analysis using pS214-Tau and GFP antibodies. Western blot analysis showed increased S214 phosphorylation and a shift in GFP-Tau phosphorylated species (upper band, arrows) upon overexpression of TBK1. In contrast, the mutant tau (Tau_null_) showed loss of S214 phosphorylation and decreased GFP-Tau phosphorylated species (upper band). B) Compared to tau alone over-expression tau and TBK1 resulted in a significant increase in S214 phosphorylation (t-test, n=3, p<0.001) C) HEK-293 cells transfected with GFP-Tau were treated with 50 μM BX795 or DMSO for 8 hours (*n*=3). Cells were harvested and subjected to western blot using pS172-TBK1, pS214-Tau and GFP antibodies. D) Addition of BX795 caused a significant decrease in pS172-TBK1 and pS214-Tau protein levels after normalization using GAPDH protein (ANOVA, n=3, ***= P<0.001, ****=P<0.0001).

### TBK1 modifies tau induced neurotoxicity in Drosophila brain

For further *in vivo* validation of our findings, we leveraged an established experimental model of AD and tauopathy, based on pan-neuronal expression of human *MAPT* in the fruit fly, *Drosophila melanogaster*. In flies, expression of either wild-type tau (tau^WT^) or mutant forms associated with FTDP-17 (tau^R406W^), leads to hyperphosphorylation and misfolding as in human disease, and triggers progressive, age-dependent neuronal dysfunction and loss (61,62). We initially examined a mutant tau^R406W^ fly strain (*Elav>tau* ^*R406W*^) (62), which is amenable to sensitive and robust detection of genetic modifiers (62); transgene expression was directed to the fly brain using a tissue-specific neuronal driver (*Elav-GAL4*). *Drosophila* has a single, well-conserved TBK1 ortholog, called *IKappaB Kinase-ε* (referred to below as *dTBK1*), encoding a protein that is 56% similar (36% identical) to human *TBK1* (*hTBK1*). Using western blots of fly head homogenates, we first confirmed that activating expression of either hTBK1 or dTBK1 in *Elav>tau*^*R406W*^ animals increases tau phosphorylation consistent with our studies in HEK-293 cells (**Figure 6A and B**). Interestingly, these manipulations also revealed an overall increase in total tau protein levels following hTBK1 or dTBK1 expression. Reciprocally, RNAi-mediated knockdown of *Drosophila* TBK1 induced relative hypophosphorylation of S214, and overall, reduced tau protein levels in *Elav>tau*^*R406W*^ animals (**Figure 6C-E**). In order to examine if genetic manipulation of dTBK1 also modulates tau-induced neurodegeneration, we aged animals 10-days and performed TUNEL staining to reveal neuronal death in the adult brain. *Elav>tau*^*R406W*^ revealed a mild degree of TUNEL-positive nuclei, as expected. However, activating expression of dTBK1 or hTBK1 significantly increased tau-mediated neuronal death, whereas knockdown of TBK1 had the opposite effect, suppressing tau neurotoxicity (**Figure 6F and G**). Importantly, either knockdown or overexpression of *TBK1* did not cause any significant neuronal death independent of tau (**Figure S3A and B**). We further confirmed genetic interactions between TBK1 and tau using an independent, complementary assay. For these experiments, Tau^WT^ was expressed in the adult *Drosophila* retina using the *Rhodopsin1-GAL4* driver, in the presence or absence RNAi targeting dTBK1, and light-induced photoreceptor polarization was measured using electroretinograms (**Figure 6H and I**). Indeed, we found that *dTBK1* knockdown is also a suppressor of Tau^WT^ neurotoxicity. We were unable to perform similar assessments of *TBK1* activation, since overexpression of either dTBK1 or hTBK1 caused electroretinogram defects independent of tau (data not shown). Overall, our studies in *Drosophila* models show that TBK1 activity modulates tau-induced neurodegeneration *in vivo*, likely via altered tau phosphorylation and levels.

**Figure 6.**
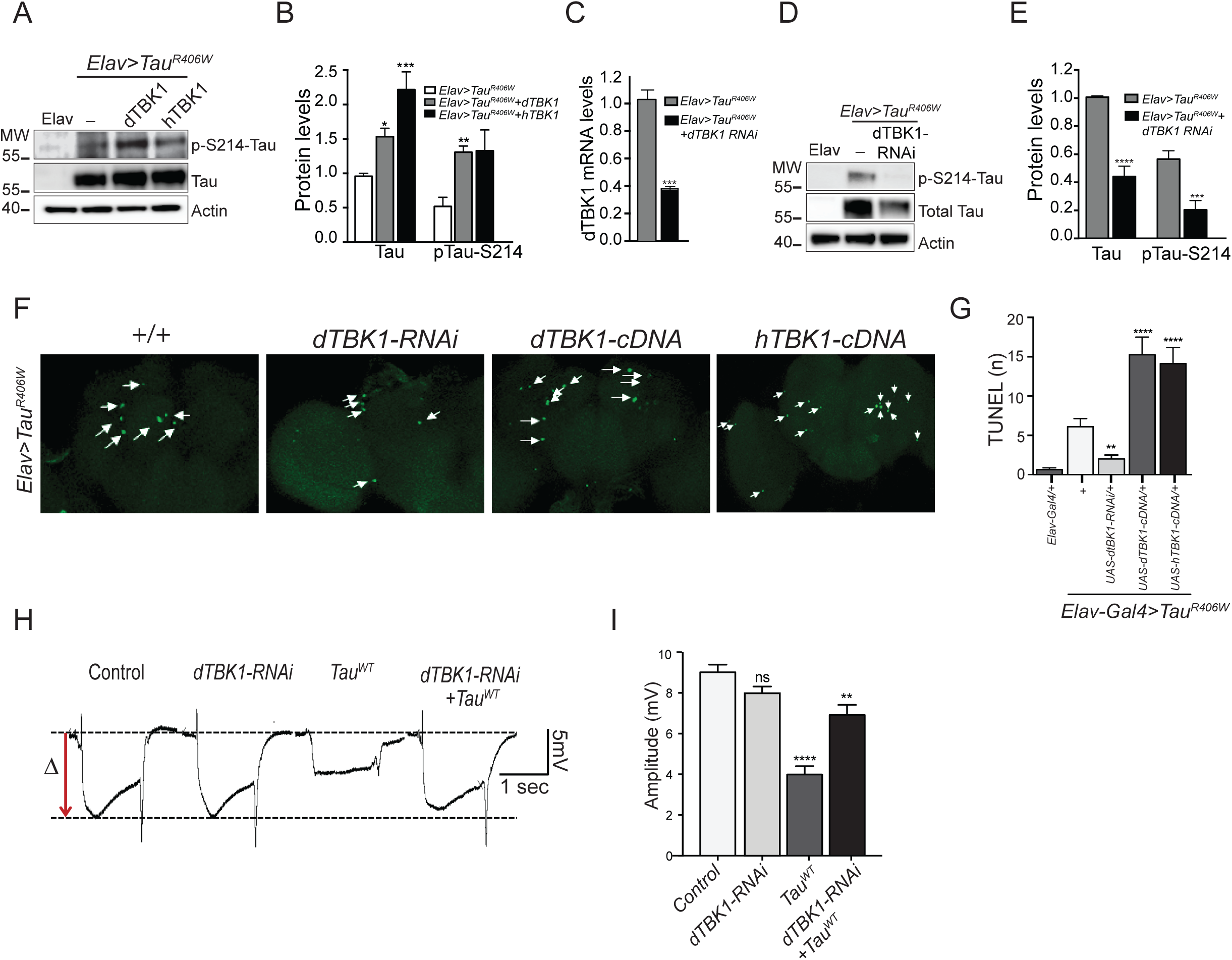
Altering TBK1 expression *in vivo* affects tau S214 phosphorylation and neurotoxicity in flies. A) Co-expression of human *TBK1* (*hTBK1*) or *Drosophila TBK1* (dTBK1) with human mutant *tau*^*R406W*^ in the adult brain (*Elav>tau*^*R406W*^ *+ UAS hTBK1-cDNA* or *Elav>tau*^*R406W*^ *+ UAS dTBK1-cDNA*) enhances tau expression and S214 phosphorylation when compared to Elav>Tau^R406W^. Western blots were prepared from fly head homogenates. Elav-Gal4 flies (Elav), without tau expression, are included as a negative control. B) Quantification of protein intensities shows increased total tau and pS214 tau upon overexpression of *hTBK1* or *dTBK1* (*n*=5 replicate experiments). Intensity levels for tau and pS214-tau were normalized to actin, and pS214-tau was further standardized relative to total tau levels. C) qPCR quantification (*n*=3) show that RNAi-mediated knockdown of *dTBK1* causes a 60% loss of mRNA levels in flies when compared to controls. D) RNAi knockdown of TBK1 suppresses tau expression and S214 phosphorylation in Elav>tau^R406W^ flies (*Elav>tau*^*R406W*^ *+ UAS dTBK1-RNAi*) when compared to controls (Elav>tau^R406W^ and Elav alone). E) Quantification of protein levels shows significant reduction in total tau and pS214 tau levels (*n*=5 replicate experiments). F) Genetic manipulation of TBK1 modifies tau-induced neurodegeneration in the adult fly brain, based on TUNEL staining in 10-day old fly brains. The same genotypes were examined as in A-E. Activating expression of hTBK1 or dTBK1 enhances whereas *dTBK1* RNAi suppressed neuronal death in *Elav>Tau*^*R406W*^ flies (*Elav>Tau*^*R406W*^ *+ UAS dTBK1-RNAi*) suppressed neuronal death when compared to controls (*Elav>Tau*^*R406W*^). G). Quantification of TUNEL-positive nuclei based on analysis of at least 10 flies for each genotype. H) dTBK1 knockdown suppresses tau^WT^ toxicity, based on retinal neurophysiology using electroretinograms. Representative traces showing light induced depolarization (Left) in the following genotypes: (i) Control (*Rh1-Gal4; +/+*), (ii) *dTBK-RNAi* (*Rh1-Gal4; UAS-dTBK-RNAi/+*), (iii) *Tau*^*WT*^ (*Rh1-Gal4; UAS-tau*^*WT*^*/+*), and (iv) *dTBK-RNAi + tau*^*WT*^ (*Rh1-Gal4; UAS-tau*^*WT*^*/UAS-dTBK-RNAi*). I) Measurement of amplitude from baseline in *n*=10 animals.

## Discussion

Tau-directed kinases play critical roles in regulating tau phosphorylation and neurofibrillary tangle formation and are considered targets for therapeutic drug development. Here, we identified TBK1 as a novel kinase that interacts with and directly phosphorylates tau. We also show increased TBK1 activation and phosphorylation of TBK1 directed sites on tau in AD and FTDP-17 (R406W). TBK1 overexpression studies in human cells and *Drosophila* neurons further confirmed the role of TBK1 activation in tau hyperphosphorylation in model systems. Finally, we show that TBK1 overexpression and knockdown in *Drosophila* can reciprocally increase and decrease tau-induced neurodegeneration.

Our previous tau interactome results from human postmortem brain tissues using co-immunoprecipitation and quantitative mass spectrometry identified increased or ‘gain-of-function’ interactions of several proteins in AD (15). Although several biological processes and targets were identified in this analysis, we focused our attention on kinases because one key feature in the cognitive progression of AD is the appearance of aberrant phosphorylation of tau. To this end, tau phosphorylation regulates its microtubule binding and stabilization, and in disease, tau hyperphosphorylation induces microtubule dissociation, which correlates with synaptic loss and neurotoxicity (7,63). Thus, identifying the kinases involved in this process, as well as developing pharmacological agents to inhibit these molecules, could be a major avenue of therapeutic interest in AD and related tauopathies. Namely, kinases such as GSK3β, CDK5 and CAMK2, previously shown to interact with or target tau (16,21), showed strong association in the AD tau interactome, highlighting the utility of our approach for mapping tau directed kinases. TBK1 was prioritized as a candidate because of its causal role in neurodegenerative diseases including FTD and ALS (29). However, other novel AD tau associated kinases identified in this study, beyond TBK1 (*i*.*e*., DAPK3 and TNIK), also warrant future investigation in their ability to phosphorylate and modify tau. For example, death-associated protein kinase-3 (DAPK3) is a Ca2+/calmodulin (CaM)-regulated serine/threonine kinases with roles in autophagy (64) and TNIK (65) has key roles in dendritic spine structure. Thus, we would hypothesize that tau is a direct substrate for these kinases and has key roles related to the respective functions of DAPK3 and TNIK in brain.

Genetic studies of *TBK1* have identified nonsense, frameshift, missense and deletion mutations in both sporadic and familial ALS, FTD and ALS/FTD, which predominately results in the loss-of-function phenotypes that reduce levels or activity of the kinase and pathological TDP-43 inclusions (45,66). Loss of TBK1 function in these diseases impacts multiple protein-protein interactions and cellular pathways including autophagy and inflammation (29,40). In contrast, TBK1 gain-of-function due to duplications are associated with rare genetic forms of glaucoma (67,68). Although we did not find differences in the steady-state levels of TBK1, our findings support an increase in TBK1 activity, and enhanced physical association with tau in AD and familial FTDP-17. Thus, TBK1 hyperactivity and mediated tau hyperphosphorylation might be a common mechanism leading to neurodegeneration observed in AD and FTLD and would align with TBK1 gain-of-function in tauopathies. However, specific TBK1 frameshift mutations that cause FTD can present with both phosphorylated TDP-43 and tau inclusions (66) indicating a possible shared mechanism for the initiation of both TDP-43 and tau pathology via TBK1 activity. Given the multiple key functions of TBK1 in cells, further investigations are suggested to resolve the direct and indirect mechanisms of TBK1 activity in neurodegenerative diseases.

Our findings support the direct ability of TBK1 to modify tau-dependent neurotoxicity. Expression of human tau in the *Drosophila* brain causes neurodegeneration, and we find that activation or reduction of TBK1 homolog enhanced or suppressed tau-mediated neurotoxicity, respectively. In future studies, it might be interesting to manipulate TBK1 in other cell types besides neurons. Indeed, studies of the mouse brain reveal TBK1 to be broadly expressed in neurons and non-neuronal cell types including microglia (69). To this end, TBK1 expression, activation, and subsequent induction of interferon signaling is increased in peripheral macrophages in response to pro-inflammatory stimuli like lipopolysaccharides (LPS) and TNF-α (31). Moreover, in mouse brain, TBK1 suppresses RIPK1 activation and downstream neuroinflammation associated with a loss of function phenotype in ALS/FTD (70). Notably, the TBK1 haploinsufficiency phenotype associated with ALS/FTD aggravates autophagy defects and early onset of motor defects, while TBK1 haploinsufficiency at later stages seems to alleviate neuroinflammation and slow disease progression in mice models (71,72). Thus, we recognize that TBK1 likely has key roles in glia and that cell type-specific phenotypes will likely differ upon global inhibition of TBK1 depending on the brain region and disease stage. This further underscore the importance of understanding mechanistically the impact of TBK1 loss-or gain-of-function in a cell-type specific manner to comprehensively define substrates and signaling pathways regulated by the kinase.

Overall, the identification of TBK1 as a novel tau modifying kinase and modifier of tau-driven neurotoxicity introduces TBK1 as a potential therapeutic target for tauopathies. TBK1 inhibitors are actively being developed for cancer and other inflammatory autoimmune diseases (73). In this study we used BX795, a well-established TBK1 inhibitor that blocks both TBK1 and IKKε (74,75). Notably, other TBK1 inhibitors including MRT67307, derivatized from BX795, suppresses TBK1 activity with much higher specificity (74). The ability of BX795 to block TBK1-directed phosphorylation *in vitro* and in cell culture lays the foundation for future studies that investigate the impact of BX795 and additional TBK1 inhibitors on tau phosphorylation and neurotoxicity in model systems.

Significant research efforts have been directed at identifying the kinases involved in tau phosphorylation as well as developing pharmacological agents to inhibit these targets. Our findings support that TBK1 is a tau-directed kinase and that increased TBK1 activity may promote tau hyperphosphorylation and neuronal loss, which makes TBK1 a potential therapeutic target for AD and related tauopathies.

## Experimental procedures

### Expression plasmids and antibodies

The following expression plasmids, pRK5-EGFP-tau (plasmid # 46904) and pWZL Neo Myr Flag TBK1 (plasmid # 20648) were purchased from Addgene. The TBK1 Open Reading Frame (ORF) was subcloned into pCDNA 3.1 expression plasmid (Emory Custom Cloning Core Facility). Site-directed mutagenesis was used to make the phospho-null tau (tau_null_) by substituting alanines for serines or threonines. Antibodies used in this study include: tau 5 (ThermoFisher, MA-12808), tau-E178 (Abcam, ab32057), pS214-tau (Abcam, ab170892), GFP (Rockland, 600-101-215), Myc (Cell Signaling, 2276), GAPDH (Abcam, ab8245), TBK1 (Abcam, ab40676), pS172-TBK1 (Abcam, ab109272), actin (Sigma, MAB1501), tau (Dako, A0224), and pS214-Tau (ThermoFisher, 44-742G).

### Human postmortem brain tissues

All human postmortem brain tissues were obtained from Emory’s Alzheimer’s Disease Research Center (ADRC) Brain Bank with prior consent. Frontal cortex brain tissue samples from AD, FTDP-17 and control cases, matched for age, gender and post-mortem interval (PMI), were selected for the study. Cases were selected based on Braak and CERAD scores that are neuropathologic measures of NFTs and amyloid plaque burden, respectively. Case traits are detailed in **Table S3**.

### Cell culture and reagents

Human embryonic kidney (HEK)-293 cells (ATCC, CRL-3216) were cultured in Dulbecco’s modified Eagle’s medium (Gibco, 11965-092) supplemented with 10% (v/v) fetal bovine serum (Gibco, 97068-091) and penicillin/streptomycin (Gibco, 15140-122) and maintained at 37°C under a humidified atmosphere of 5% (v/v) CO_2_. For transient transfection, cells grown to 80–90% confluency were transfected with expression plasmids using JetPrime reagent (Polyplus, 114-07) according to the manufacturer’s instructions. A TBK1 inhibitor BX795 (ENZ-CHM189-0005), was added at the reported concentrations 12 hrs. post transfection and further incubated for 8 hrs. Subsequently, cells were rinsed with cold 1x phosphate buffered saline (PBS) and harvested in NP-40 lysis buffer (25 mM Tris-HCl (pH 7.5), 150 mM NaCl, 1 mM EDTA, 1% NP-40, 5% glycerol, plus HALT protease inhibitor cocktail, 87786).

### Brain tissue homogenization

Human postmortem brain tissues were homogenized as previously described(15). Briefly, tissues were homogenized in lysis buffer (25 mM Tris-HCl (pH 7.5), 150 mM NaCl, 1 mM EDTA, 1% NP-40, 5% Glycerol, and HALT protease inhibitor cocktail) using a bullet blender and the protein lysate was cleared by centrifugation at 10,000 x g for 15 min, followed by bicinchoninic acid (BCA) analysis (Pierce, 23225) to determine protein concentration.

### Western blotting

Protein samples were boiled in Laemmli sample buffer at 98°C for 5 min, resolved on Bolt 4-12% Bis-Tris gels (Thermo Fisher Scientific, cat. # NW04120BOX), followed by transfer to nitrocellulose membrane using iBlot 2 dry blotting system (ThermoFisher Scientific, IB21001). Membranes were incubated with StartingBlock buffer (ThermoFisher, 37543) for 30 min followed by overnight incubation in primary antibodies at 4°C. Membranes were washed with 1×Tris-buffered saline containing 0.1% Tween 20 (TBS-T) and incubated with fluorophore-conjugated secondary antibodies (AlexaFluor-680 or AlexaFluor-800) for 1 hr. at room temperature. Then, membranes were washed three times with TBS-T and images were captured using an Odyssey Infrared Imaging System (LI-COR Biosciences). For *Drosophila*, 10-day old fly heads lysates were isolated and prepared in Laemmli sample buffer for western blot analyses as previously described(15).

### Co-immunoprecipitation assay (co-IP)

Co-IP was done using 1 mg of protein from brain or HEK-293 cell lysates. Samples were first pre-cleared by incubating with Protein A-Sepharose conjugated beads (Invitrogen, 101041) for 1 hr. at 4 °C. Lysates were then incubated with 5 μg of primary antibodies (Tau-5, GFP, Myc) overnight at 4 °C. Purified Mouse IgG2a K isotype (BD pharmingen, 550339) was used as an isotype-matched negative control. After overnight incubation, 50 μL of DynaBeads Protein G suspension (Invitrogen, 10003D) was incubated with lysates for 1 hr. Beads were then washed three times using wash buffer (50 mM Tris-HCl, pH 8.0, 150 mM NaCl and 0.1% NP-40), rinsed three times using 1x PBS and boiled in 40 μl Laemmli sample loading buffer for western blot analysis as described above.

### *In vitro* kinase assay

*In vitro* kinase assays were performed by incubating 0.5 μg active TBK1 (SignalChem, T02-10G-10) with 1 μg tau 441 purified recombinant proteins (SignalChem, T08-55BN-20) in kinase assay buffer (25 mM MOPS, pH 7.2, 12.5 mM β-glycerol-phosphate, 25 mM MgCl_2_, 5 mM EGTA, 2 mM EDTA and 0.25 mM DTT) plus 100 μM ATP for 30 min at 37°C (SignalChem, K01-09). TBK1 kinase activity was inhibited by adding 100 μM or 200 μM of BX795 (ENZ-CHM189-0005) (59,60). The kinase assay was quenched by boiling samples at 98°C in Laemmli sample buffer (BioRad, 161-0737) for 5 min prior to western blot analysis.

### Mass spectrometry and tau phosphopeptide quantification

For mass spectrometry analyses the TBK1 kinase activity assay reaction was resuspended in 100 μL 50 μM NH_4_HCO_3_ buffer and subjected to standard tryptic digestion and MS analysis as described (76,77). MS raw files were searched using MaxQuant (78) (version 1.6.3.4) search platform for identification and quantification of tau phosphorylation sites (**Table S2**). Human postmortem brain tau IP-MS raw files(15) were searched using the MaxQuant search engine to identify and quantify tau phosphopeptides as previously described (79) (**Table S1**).

### Immunohistochemistry

Paraffin embedded human postmortem brain sections (5 μm) were deparaffinized in Histo-clear (National Diagnostics) and rehydrated in ethanol. Antigen retrieval was performed in ddH2O by steam heat for 30 minutes. Endogenous peroxidase activity was quenched using hydrogen peroxide and washed 3x in TBS-T. Tissues were blocked with serum-free protein block (Dako) for 1 hour. Primary antibodies were diluted in antibody diluent (Dako) and applied for 45 minutes at room temp. Tissue sections were subsequently washed 3x in TBS-T and HRP-conjugated secondaries (Dako) were applied for 30 minutes at room temperature. 3,3’-diaminobenzidine (DAB) was used for visualization of peroxidase labeling. Sections were counterstained with Gill’s hematoxylin and blued in Scott’s tap water substitute.

### Gene Ontology (GO) Enrichment and Hierarchical Clustering Analysis

Functional enrichment of differentially expressed proteins was determined using the GO-Elite (v1.2.5) python package as previously described (76,80). Briefly, GO-Elite Hs (human) databases were downloaded on or after June 2016. The set of all proteins identified in the proteomic analyses was used as the background. Z score and one tailed Fisher’s exact test (Benjamini-Hochberg FDR corrected) was used to assess the significance of the Z score. Peptides were collapsed to one gene symbol and each gene symbol was equally weighted regardless of the number of matching peptides. Z score cut-off of 1.96, P value cut off 0.01 and a minimum of 5 genes per ontology were used as filters prior to pruning the ontologies. Clustering analysis on differentially abundant phospho-modified peptides was performed with the R NMF package in Microsoft R open v3.3. Age, sex and PMI-regressed log_2_ abundances were converted to Z scores (mean centered abundance, fold of standard deviation) and clustered with Euclidian distance metric, complete method of the hclust function called from NMF package aheatmap function.

### *Drosophila* genetics

Flies were raised on molasses-based food grown at 20°C until eclosion and then placed at 25°C for 10 days. Fly strains were obtained from the Bloomington *Drosophila* Stock Center and from the FlyORF. The Drosophila ortholog of Human TBK1 is *IKK-epsilon* (IKKε or ik2) and is referred to in this manuscript as dTBK1. The Full genotypes of flies used are as follow. UAS-*dTBK1*-RNAi(P{TRIP.GL00160}attP2, BDSC_35266; Flybase ID: FBal0262637), UAS-*dTBK1*-cDNA (M{UAS-IKKε.ORF.3xHA}ZH-86Fb, FlyORF_F001016; Flybase ID: FBst0500628), and UAS-*hTBK1*-cDNA (PBac{UAS-hTBK1.HA}, FlyORF_77971; Flybase ID: FBst0077971). *Elav*^*c155*^*-Gal4* flies were used to drive expression of *UAS* lines pan-neuronally in the adult fly brain and the tauopathy model used in this manuscript expresses human mutant Tau (UAS-Tau^R406W^) and human wild type Tau (UAS-Tau^WT^) (62).

### Quantitative Real-Time PCR (qRT-PCR)

Total RNA was extracted from approximately 100 adult fly heads (for each genotype), equally divided between males and females. Frozen fly heads were homogenized in Trizol (Invitrogen, 15596026), treated with DNaseI (Promega, M6101), and total RNA was extracted using the RNeasy Micro Kit (QIAGEN, 74004). Following reverse transcription using the SuperScript III First-Strand Synthesis System (Invitrogen), qRT-PCR was performed using iQ SYBR Green Supermix (Bio-Rad) in a CFX96 Touch Real-Time PCR Detection System (Bio-Rad) with standard cycling parameters. Each reaction was performed in triplicate. RpL32 was used as a control for normalization of each sample to calculate delta cycle threshold (DCt) values.

### *Drosophila* terminal deoxynucleotidyl transferase dUTP nick end labeling (TUNEL) Assay

TUNEL staining was performed using the FragEL DNA Fragmentation Detection Kit from Calbiochem (EMD Millipore). 10-day old adult fly brains were dissected in PBST and then fixed in 4% paraformaldehyde for 20 minutes on ice. Brains were washed with 1x phosphate-buffered saline with 0.1% Tween 20 (PBS-T) 4 times for 15 minutes and then blocked overnight with 5% normal goat serum (NGS) in PBS-T at 4°C. Brains were then incubated in 100 mM Sodium Citrate with 10% Triton x-100 for 30 minutes at 65 °C and then washed 3 times with PBST at room temperature. Brains were equilibrated for 15 minutes at room temperature, and then incubated in a 1:9 mixture of Terminal deoxynucleotidyl transferase (Tdt) enzyme and Tdt Labeling Buffer for 2-3 hours at 37 °C. Brains were washed 4 times for 15 minutes at room temperature and then mounted on slides with Vectashield (with DAPI). Images were acquired using Z stack with a 2.00 step at 20x using a SP8 Leica confocal microscope and apoptotic neurons were counted.

### *Drosophila* electroretinograms (ERG)

ERG recordings were performed as previously described (81). In brief, 10-day old adult flies were anesthetized and glued to a glass slide, with electrodes placed on the corneal surface and the thorax. Flies were maintained in the dark for at least 1 min prior to stimulation using a train of alternating light/dark pulses. Retinal responses were recorded and analyzed using LabChart software (ADInstruments). At least eight flies were examined for each genotype.

## Data availability

The mass spectrometry proteomics data from the Tau co-IPs from human postmortem brain samples have been deposited to Synapse. http://synapse.org/TauBrainIP DOI: 10.7303/syn22150694

## Acknowledgements

We would like to thank members of the Seyfried lab including Dr. Sruti Rayaprolu, Sean Kundinger and Dr. Pritha Bagchi for their feedback.

## Funding and additional information

Support for this research was provided by funding from the National Institute on Aging (R01AG050631, R01AG053960, R01AG061800, RF1AG057471, RF1AG057470, R01AG061800, R01AG057339), the Accelerating Medicine Partnership (AMP) for AD (U01AG046161 and U01AG061357) and the Emory Alzheimer’s Disease Research Center (P50AG025688). S.A.O. was supported by a Postdoctoral Enrichment Program Award from the Burroughs Welcome Fund (BWF-1017399) and an Alzheimer’s Association fellowship (AARFD-16-442630).

## Conflict of interest statement

None declared.

## Supporting Information

**Supplemental Figure S1.**
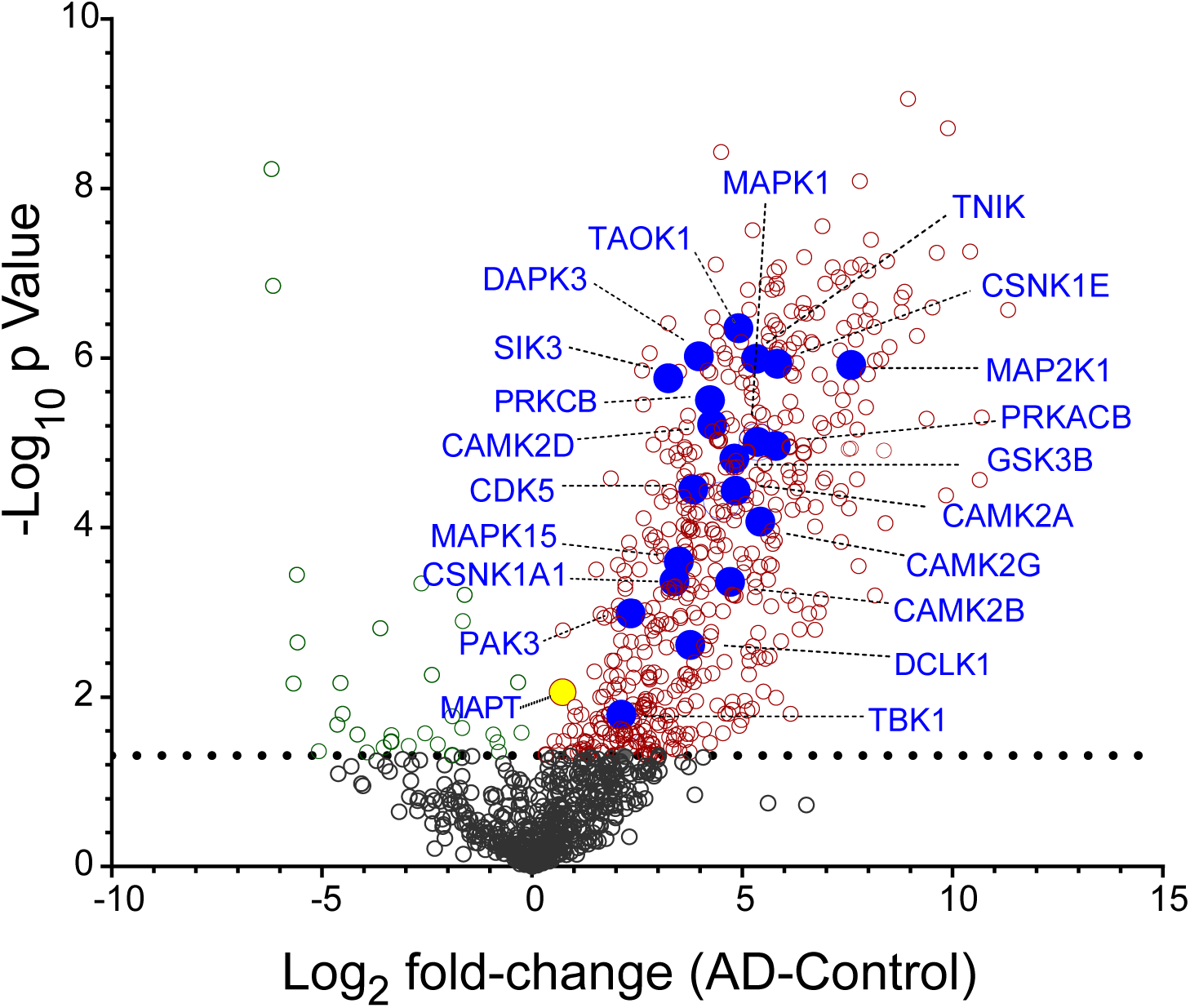
Tau interactome in AD brain. MS analysis of tau immunoprecipitates from AD (*n*=4) and control (*n*=4) postmortem brain homogenates identified 1,065 proteins. A volcano plot with log_2_-fold-change (AD-Control) on the x-axis against −log_10_ *P* value (y-axis) displays differential tau interaction. Proteins above the dashed line are statistically significant by t-test (p<0.05). Proteins with at least 1.5-fold increased interaction in AD vs control are depicted in red open circles (n=513) whereas proteins with at least 1.5-fold decreased interaction in AD are depicted in green open circles (n=29). All tau interacting kinases are depicted in purple circles and labeled.

**Supplemental Figure S2.**
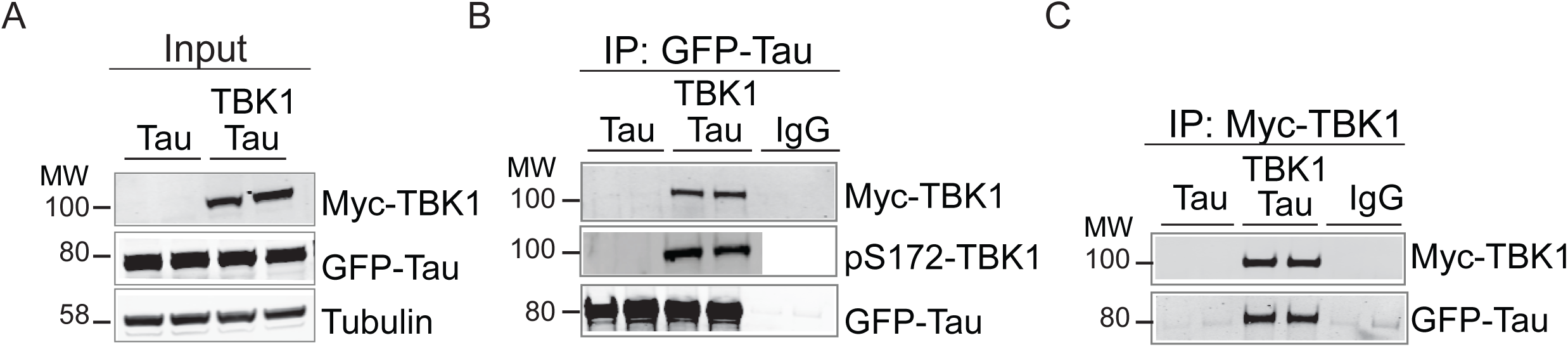
Tau and TBK1 interactions in HEK-293 cells. A) Western blotting using Myc and GFP antibodies shows expression of both recombinant tau-GFP and myc-TBK1 in co-transfected cells. Immunoprecipitation of tau using GFP antibody followed by western blotting using Myc and pS172- TBK1 antibodies showed positive interactions of tau with TBK1 and activated TBK1 (pS172-TBK1), respectively, in HEK-293 cells. C) Reciprocal IP using Myc antibody and western blotting for GFP showed Tau and TBK1 interaction in HEK-293 cells. HEK-293 lysate expressing tau alone and bead only controls were included as negative controls. Tubulin is used as a loading control.

**Supplemental Figure S3.**
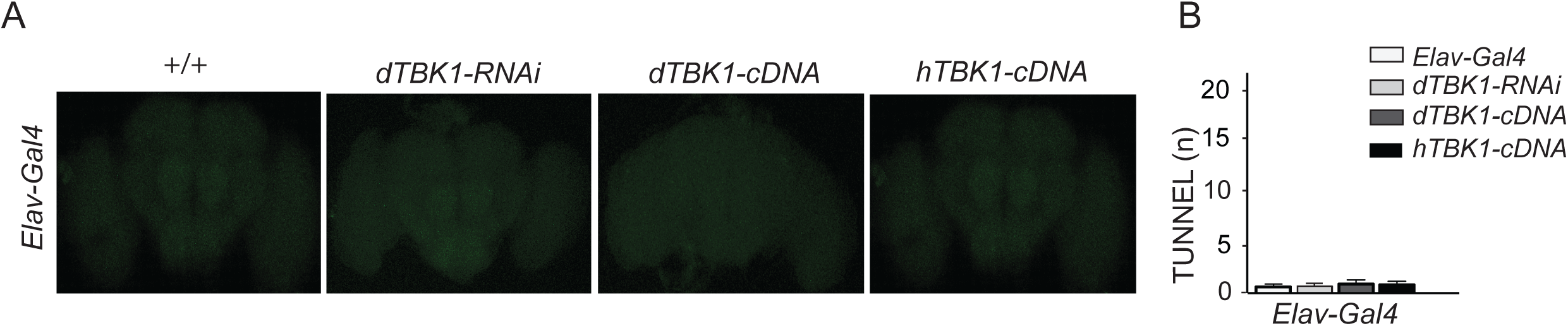
Altering TBK1 expression in the absence of tau does not cause toxicity. A) Pan-neuronal knockdown (*Elav>dTBK1-RNAi*) or over-expression (*Elav>hTBK1-cDNA* or *Elav>dTBK1- cDNA*) of *TBK1* in 10-day old adult fly brains does not cause any significant neuronal death when compared to controls (*Elav*) as show by TUNEL staining. B) Quantification of TUNEL-positive nuclei based on analysis of at least 10 flies for each genotype.

## Notes

### Competing Interest Statement

The authors have declared no competing interest.

